# *Borrelia burgdorferi* Secretes c-di-AMP as an Extracellular Pathogen-Associated Molecular Pattern to Elicit Type I Interferon Responses in Mammalian Hosts

**DOI:** 10.1101/2024.08.13.607721

**Authors:** Raj Priya, Meiping Ye, Sajith Raghunanadanan, Qiang Liu, Wei Li, Qigui Yu, Yongliang Lou, Herman O. Sintim, X. Frank Yang

## Abstract

*Borrelia burgdorferi* (*B. burgdorferi*), an extracellular spirochetal pathogen, elicits a type-I interferon (IFN-I) response that contributes to the pathology of Lyme disease, including the development and severity of Lyme arthritis. However, the specific Pathogen-Associated Molecular Patterns (PAMPs) of *B. burgdorferi* responsible for triggering the IFN-I response are not well understood. Previous studies have identified an unknown, nuclease-resistant component in *B. burgdorferi* culture supernatants that significantly stimulates the IFN-I response, but its identity remains unknown. In this study, we reveal that *B. burgdorferi* secretes cyclic-di-adenosine monophosphate (c-di-AMP) as a key extracellular PAMP, inducing the host IFN-I response in macrophages. Using genetically manipulated *B. burgdorferi* strains, we demonstrate a requirement of c-di-AMP for stimulating IFN-I response by macrophages *ex vivo*. Additionally, infecting mice with *B. burgdorferi* alongside exogenous c-di-AMP resulted in a markedly increased IFN-I response in mouse tissues. Furthermore, inactivation or inhibition of the host STING signaling pathway significantly reduced the IFN-I response, indicating that c-di-AMP-induced IFN-I production is STING-dependent. Our findings identify c-di-AMP as a crucial PAMP secreted by *B. burgdorferi* to elicit the host IFN-I response via activation of STING signaling pathway, suggesting that targeting c-di-AMP production could represent a novel therapeutic strategy against Lyme arthritis.

**SUMMARY:** *Borrelia burgdorferi*, the bacteria that causes Lyme disease, induces a robust host immune response, including the production of type-I interferon (IFN-I). While this response helps combat the infection, it also contributes to complications such as Lyme arthritis. Our research aimed to identify the specific bacterial component that triggers the IFN-I response. We discovered that *Borrelia burgdorferi* releases a second messenger molecule, cyclic-di-adenosine monophosphate (c-di-AMP), which is recognized by host immune cells and subsequently triggers IFN-I production. This finding is significant as it advances our understanding of Lyme disease pathogenesis and offers a new strategy to tackle Lyme disease by targeting the production of c-di-AMP, in which we may be able to reduce the severity of the disease and mitigate long-term tissue damage.

**One sentence summary:** *Borrelia burgdorferi* c-di-AMP induces Type I IFN response

## INTRODUCTION

Lyme disease is the most commonly reported arthropod-borne illness in the US, Europe, and Asia (Radolf et al., 2021, Burgdorfer et al., 1982, Lochhead et al., 2021). It is caused by genospecies of the *Borrelia burgdorferi* sensu lato complex, spirochetal bacteria with dual membranes that lack lipopolysaccharides (LPS) commonly observed in typical Gram-negative bacteria. *B. burgdorferi* is transmitted to humans via *Ixodies* tick bites. If left untreated, it results in a multi-stage and multi-organ infection, starting with a local skin lesion (erythema migrans) and progressing to disseminated systemic infection, leading to Lyme arthritis, carditis, and neurological disorder. The tissue pathology of the disease is primarily due to host inflammatory reactions in response to the infection of *B. burgdorferi*. Abundant lipoproteins on the surface of *B. burgdorferi* are known as the major pathogen-associated molecular patterns (PAMPs) recognized by the host pattern recognition receptor (PRR), Toll-like receptor 2 (TLR2), leading to potent proinflammatory cytokines and innate immune response (Bockenstedt *et al*., 2021). In addition, *B. burgdorferi* induces type I interferon (IFN-I) response typically associated with viral and intracellular bacterial infections (McNab *et al*., 2015). It has been reported that *B. burgdorferi* strains associated with disseminated Lyme disease in humans and mice induce high levels of IFN-I response (Petzke *et al*., 2016). More importantly, IFN-I is critical for the development of subacute Lyme arthritis in infected mice (Miller *et al*., 2008). Additionally, recent reports have shown a link between persistent symptoms following Lyme neuroborreliosis and elevated IFN-I levels in the blood of Lyme disease patients (Hernández *et al*., 2023, Jacek *et al*., 2013). These findings underline the significance of the IFN-I response in multiple aspects of Lyme disease pathogenesis. However, how *B. burgdorferi* induces IFN-I response in the host has not been fully elucidated.

Induction of IFN-I response to virus and bacteria is mediated through either TLR-dependent or independent signaling pathways. The TLR-dependent signaling pathway can be further categorized into the Myd88- or TRIF-II-dependent. In this regard, *B. burgdorferi*-induced transcription of IFN-responsive genes appear to be MyD88 and TRIF-independent (Miller *et al*., 2008, Miller *et al*., 2010). Cytosolic TLR-independent signaling pathways primarily include NOD-like receptors (NLRs), retinoic acid-inducible gene I (RIG-I)-like receptors (RLRs), and the cyclic guanosine monophosphate (GMP)-adenosine monophosphate (AMP) (cGAMP) synthase (cGAS) and stimulator of interferon genes (STING) (together called cGAS-STING) signaling pathways. It was reported that the NLRs and RLRs receptors are not required for the induction of IFN-I response by *B. burgdorferi* (Miller *et al*., 2008, Farris *et al*., 2023). On the other hand, recent seminal work by Farris *et al*. demonstrates that *B. burgdorferi* activates the cGAS-STING signaling pathway, which significantly contributes to the production of IFN-I (Farris *et al*., 2023).

The cGAS-STING signaling pathway is an important part of the host’s innate immune system that protects the host from external pathogens and internal damage. The classical cGAS-STING signaling pathway activation occurs upon the cGAS protein interacting with either microbial DNA or cytosolic damaged-associated host DNA, catalyzing the production of 2′3′-cGAMP from ATP and GTP, which subsequently binds and activates endoplasmic reticulum (ER)-localized STING. Activated STING translocate from the ER through the Golgi apparatus to perinuclear vesicles. During this process, STING recruits and activates the kinase TBK1, which subsequently phosphorylates the transcription factor IRF3. Phosphorylated IRF3 then dimerizes and translocates to the nucleus, where it induces IFN-I response (Yum *et al*., 2021). It has well recognized that the cGAS-STING signaling pathway is activated during infection by intracellular pathogens, as well as some extracellular pathogens (Chen *et al*., 2016, Liu *et al*., 2022, Monroe *et al*., 2010, Andrade *et al*., 2016). The finding that *B. burgdorferi* activates the cGAS-STING signaling pathway to induce an IFN-I response further reinforced this notion (Farris *et al*., 2023). Despite of these findings, the specific PAMP of *B. burgdorferi* recognized by host cells to elicit an IFN-I response remains elusive.

Common PAMPs in bacteria triggering the IFN-I response include DNA, RNA, lipoproteins, peptidoglycans, or lipopolysaccharides (LPS) (Akira *et al*., 2006, Kawai & Akira, 2010). *B. burgdorferi* lacks LPS (Takayama *et al*., 1987). Previous reports showed that *B. burgdorferi* induces an IFN-I response independently of lipoproteins and peptidoglycans (Miller *et al*., 2010, Miller *et al*., 2008). The role of DNA as a PAMP of *B. burgdorferi* to induce the IFN-I response has been controversial. Petzke et al. and Farris et al. suggested that DNA is important for the induction of the IFN-I response (Petzke *et al*., 2009, Farris *et al*., 2023), while Miller et al. demonstrated that DNA plays a dispensable role (Miller *et al*., 2008, Miller *et al*., 2010). However, *B. burgdorferi* RNA can induce an IFN-I response. Remarkably, Miller et al., also found that *B. burgdorferi* culture supernatants can strongly stimulate the expression of IFN-I-responsive genes. This unknown PAMP secreted by *B. burgdorferi* appears to be DNase- and RNAse-resistant, but its exact nature remains unidentified (Miller *et al*., 2010).

In recent years, bacterial second messengers, cyclic dinucleotides, in particular, c-di-AMP, have been recognized as important PAMPs that stimulate host IFN-I response (McNab *et al*., 2015, Wright & Bai, 2023, Woodward *et al*., 2010, Dey *et al*., 2015, Moretti *et al*., 2017). c-di-AMP is found mostly in Gram-positive bacteria (Stülke & Krüger, 2020, Witte *et al*., 2008). It is synthesized by diadenylate cyclase (DAC) and degraded by phosphodiesterase (PDE). c-di-AMP plays important roles in many bacterial processes including antibiotic resistance, biofilm formation, cell wall synthesis, peptidoglycan homeostasis, DNA repair, potassium homeostasis, and osmotic stress response (Stülke & Krüger, 2020). In addition, it can function as a cross-kingdom signal between bacteria and eukaryotic hosts by directly binding to and activating the ER-associated STING, leading to a robust IFN-I response (Woodward *et al*., 2010, Barker *et al*., 2013, Dey *et al*., 2015, Gries *et al*., 2016, Wright & Bai, 2023, Zaver & Woodward, 2020, Andrade *et al*., 2016). Furthermore, unlike classic PAMPs such as LPS shared by both live and dead pathogens, c-di-AMP has been regarded as a vita-PAMP (viability-associated PAMP) of live pathogens, by triggering more robust immune responses compared to dead pathogens. (Moretti *et al*., 2017).

*B. burgdorferi*, a Gram-negative like spirochete, is distinct in possessing a diadenylate cyclase (CdaA) and a phosphodiesterase (DhhP), enzymes typically found in Gram-positive bacteria (Ye *et al*., 2014, Savage *et al*., 2015). Previously, we showed that the *dhhP* mutant, which produced constitutively high intracellular c-di-AMP, failed to grow *in vitro* and is incapable of infecting mice (Ye *et al*., 2014). However, the function of c-di-AMP in the physiology and pathogenesis of *B. burgdorferi* remains poorly understood. Given the fact that *B. burgdorferi* activates cGAS-STING signaling to induce IFN-I response, we postulate that *B. burgdorferi* may use c-di-AMP as a PAMP to induce IFN-I response. In this study, we tested this hypothesis by employing *B. burgdorferi* mutants producing varying levels of c-di-AMP and utilizing macrophages with either functional or non-functional STING to elucidate the impact on IFN-I response. We found that *B. burgdorferi* is capable of secreting c-di-AMP, and the extracellularly secreted c-di-AMP contributes significantly to the activation of STING signaling resulting in the induction of IFN-I response. Thus, c-di-AMP appears to be the key component in the spent culture supernatant observed previously (Miller *et al*., 2010) that stimulates the IFN-I response and functions as an extracellular vita-PAMP of *B. burgdorferi*, a Gram-negative-like pathogen.

## RESULTS

### *B. burgdorferi* produces and secretes c-di-AMP

*B. burgdorferi* harbors a *cdaA* and a *dhhP* gene, which encode for the synthesis and degradation of c-di-AMP, respectively (Ye *et al*., 2014, Savage *et al*., 2015). We previously reported that the *dhhP* mutant constitutively produces a high level of c-di-AMP (Ye et al., 2014). To determine the contribution of CdaA to c-di-AMP levels in *B. burgdorferi*, a *cdaA* mutant was constructed using the allele exchange method as previously described (**supplemental Fig. S1**) (Zhang *et al*., 2020). Unlike the *dhhP* mutant, the cdaA mutant had no growth defect *in vitro* (Ye *et al*., 2014). The intracellular levels of c-di-AMP in wild-type, *cdaA* mutant, and *dhhP* mutants were determined using a highly sensitive competitive ELISA assay (Underwood *et al*., 2014). The results showed that wild-type *B. burgdorferi* (5A4NP1) produces a low but significant level of c-di-AMP (**Fig. 1A**). As expected, the *dhhP* mutant significantly produced increased c-di-AMP levels by 28-fold as compared to the wild-type strain (**Fig. 1A**). In contrast, the *cdaA* mutant had dramatically lower c-di-AMP levels (3.5-fold less than that of the wild-type strain). This result further confirms that CdaA and DhhP are the c-di-AMP cyclase and phosphodiesterase of *B. burgdorferi*, respectively.

**Fig 1.**
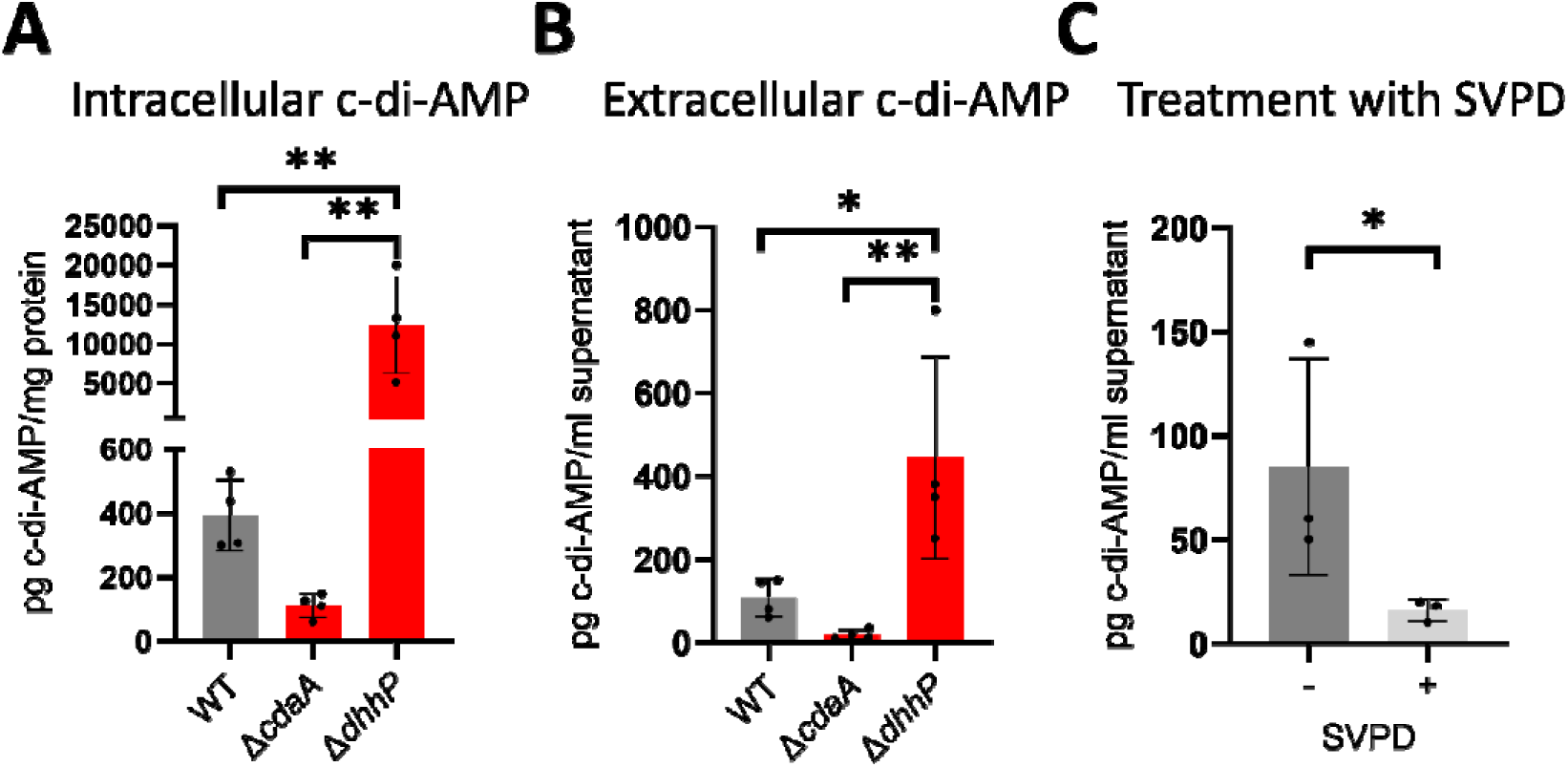
*B. burgdorferi* produces and secretes c-di-AMP *in vitro*. Wild-type (WT) *B. burgdorferi* 5A4NP1, Δ*cdaA*, and Δ*dhhP* were cultured until the late logarithmic phase. Culture pellets and supernatants were collected separately. The levels of **(A)** intracellular (in bacterial cells) and **(B)** extracellular (in culture supernatants) c-di-AMP levels were analyzed using c-di-AMP ELISA assays. **(C)** The culture supernatants of wild-type *B. burgdorferi* were treated with either water (control) or phosphodiesterase (SVPD) before c-di-AMP ELISA assays. The data represent the mean ± SD from at least three experiments (n = 3). Statistical significance was assessed using the one-way ANOVA for **(A)** and **(B)** and two-tailed Student’s t-test for **(C)** with \**p* < 0.05; \*\**p* < 0.01.

To investigate whether *B. burgdorferi* is capable of secreting c-di-AMP, supernatants from wild-type and mutant strains of *B. burgdorferi* cultures were collected and subjected to ELISA assays. We first observed that the BSK-II growth medium used for culturing *B. burgdorferi*, a nutrient-rich medium containing a high concentration of BSA and rabbit serum, interfered with the c-di-AMP ELISA assay. Thus, the culture supernatants were first diluted in DMEM (5-fold dilution) before the measurement. As shown in **Fig. 1B**, c-di-AMP was readily detectable in the spent culture supernatant of wild-type *B. burgdorferi*, with a 4-fold increase in the Δ*dhhP* strain and a 5.6-fold reduction in the Δ*cdaA* strain. To further confirm that the signals detected in the culture supernatants were c-di-AMP-specific, the culture supernatants were treated with snake venom phosphodiesterase (SVPD), an enzyme known to degrade bacterial c-di-AMP and then utilized for ELISA assay (Woodward *et al*., 2010, Barker *et al*., 2013). Pre-treatment of the culture supernatant with SVPD significantly reduced the c-di-AMP levels (by 5.3-fold) (**Fig. 1C**). These results suggest that *B. burgdorferi* is capable of secreting c-di-AMP during *in vitro* growth.

### c-di-AMP plays a key role in *B. burgdorferi*-induced IFN-I response

To determine whether c-di-AMP is a PAMP of *B. burgdorferi* for IFN-I induction, murine macrophage cell lines (RAW264.7) were infected with either wild-type or mutant strains of *B. burgdorferi* at an MOI of 10 for 24 hours. The level of IFN-β in the macrophage supernatant in response to *B. burgdorferi* infection was determined using the ELISA. As expected, wild-type *B. burgdorferi* induced IFN-β production in macrophages (**Fig. 2A**). However, the *cdaA* mutant induced significantly lower levels of IFN-β (2.2-fold less than wild-type *B. burgdorferi*), whereas the *dhhP* mutant, which produced high levels of c-di-AMP, induced higher levels of IFN-β (2-fold more than wild-type strain) (**Fig. 2A**). This finding was further confirmed using bone marrow-derived macrophages (BMDMs) isolated from C3H/HeN mice (**Fig. 2D**). These results suggest that c-di-AMP contributes significantly to the *B. burgdorferi*-induced IFN-I response.

**Fig. 2.**
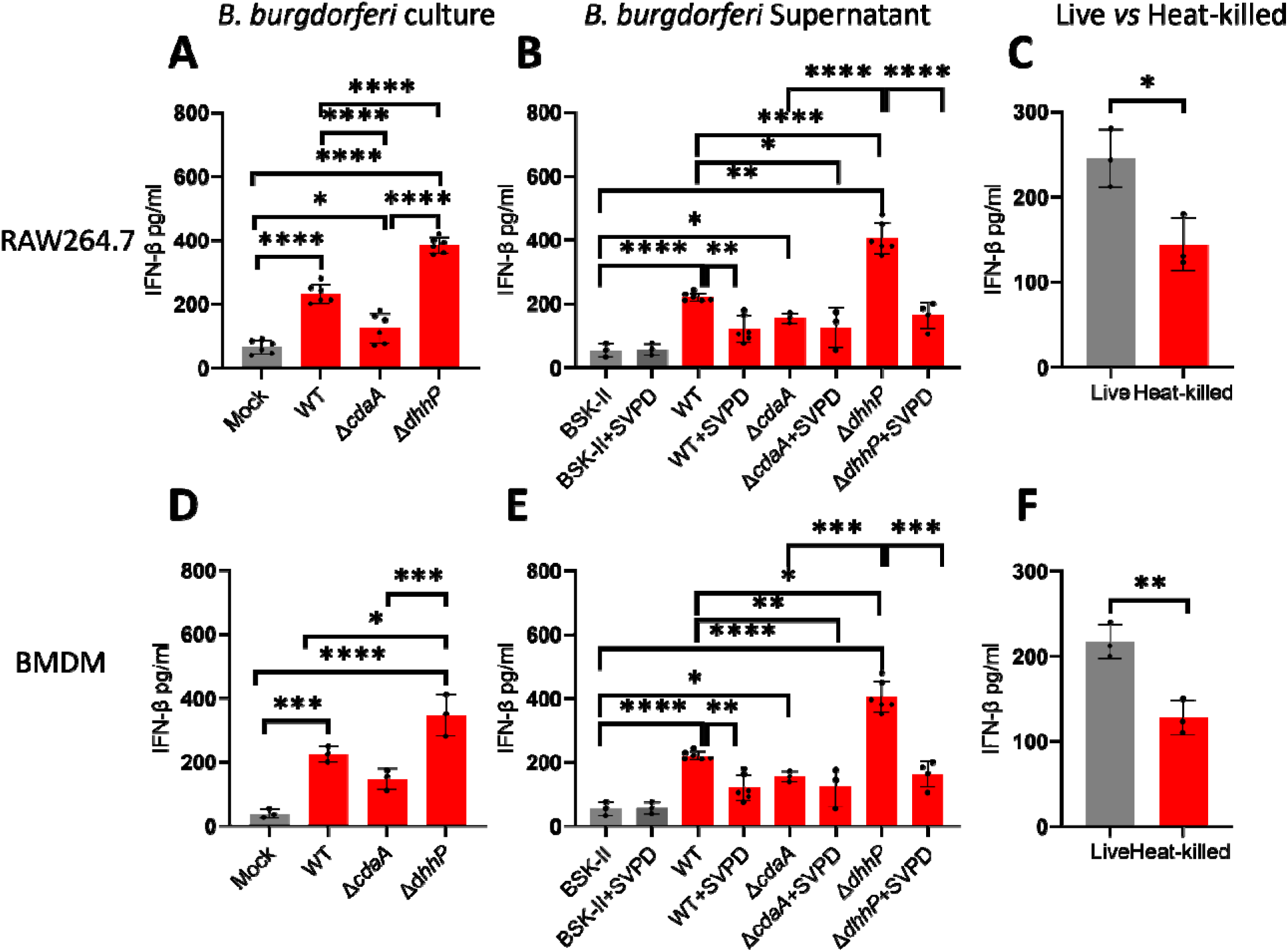
c-di-AMP is important for the induction of IFN-β production by macrophages following infection with *B. burgdorferi*. Wild-type (WT) *B. burgdorferi* 5A4NP1, Δ*cdaA*, and Δ*dhhP* were cultured until the late logarithmic phase. *B. burgdorferi* culture pellets and spent supernatants were collected separately. RAW (**A**) and BMDMs (**D**) were infected with either *B. burgdorferi* at a MOI of 10 or treated with diluted spent supernatants either pre-treated with water (control) or phosphodiesterase (SVPD) (**B, E**). Live or heat-killed *B. burgdorferi* cultures were also used to infect RAW cells (**C**) or BMDMs (**F**) at a MOI of 10. In all experiments, macrophage supernatants were collected after 24 hours of treatment and subjected to ELISA assays for IFN-β levels. The data represent the mean ± SD of at least three experiments (n = 3-6). Statistical significance was assessed using the one-way ANOVA for (**A**), (**B**), (**D**), and (**E**), and two-tailed student’s t-test for (**C**) and (**F**) with \**p* < 0.05; \*\**p* < 0.01; \*\*\**p* < 0.001.

Previously, Miller et al., demonstrated that the supernatant of *B. burgdorferi* culture significantly stimulates IFN-I production by macrophages (Miller *et al*., 2010). To confirm this, diluted *B. burgdorferi* culture supernatants were incubated with RAW macrophages for 24 hours and the levels of secreted IFN-β by macrophages were measured. As shown previously, the supernatant from wild-type *B. burgdorferi* culture induced IFN-β by macrophages (**Fig. 2B**). Importantly, elevated IFN-β induction was observed (1.8-fold higher than the wild-type group) when macrophages were treated with the supernatant from the Δ*dhhP* culture and decreased IFN-β induction was observed (1.4-fold lower than wild-type group) when treated with the supernatant from the Δ*cdaA* culture (**Fig. 2B**). Notably, treatment of supernatants of *B. burgdorferi* cultures with SVPD before exposure to macrophages reduced the level of IFN-β production (**Fig. 2B**). These findings were further confirmed by using bone marrow-derived macrophages (BMDMs) isolated from C3H/HeN mice (**Fig. 2E**). These results indicate that c-di-AMP, secreted by *B. burgdorferi*, is a crucial component of the *B. burgdorferi* culture supernatant that triggers a robust IFN-I response in macrophages.

c-di-AMP has been recognized as a vita-PAMP as it is a signature of the viable intracellular pathogen (Moretti *et al*., 2017), which elicits stronger immune responses than classic PAMPs present in dead bacteria. Consistent with this, macrophages infected with live *B. burgdorferi* exhibited significantly higher IFN-β production than heat-killed bacteria (by 1.7-fold in RAW cell lines or BMDMs) (**Fig. 2C and 2F**). Thus, c-di-AMP is a vita-PAMP of the extracellular, Gram-negative bacteria like *B. burgdorferi*.

### Exogenous c-di-AMP enhances *B. burgdorferi-*induced IFN-I response *in vitro*

To further confirm that c-di-AMP is the PAMP of *B. burgdorferi* that stimulates IFN-I response, exogenous c-di-AMP (1 µg/ml) was added to the co-culture of RAW264.7 macrophages and *B. burgdorferi*. After 24 hours of infection, levels of secreted IFN-β protein and levels of *Ifnb1*, *Cxcl10*, and *Ifit1* mRNAs in macrophages were determined. The result showed that adding exogenous c-di-AMP to the *B. burgdorferi*-macrophage co-culture significantly enhanced the level of IFN-β production by macrophages compared to the *B. burgdorferi* infection alone (**Fig. 3A**). Similarly, increases in the levels of *Ifnb1*, *Cxcl10*, and *Ifit1* mRNAs were observed when adding exogenous c-di-AMP to the co-culture (**Fig. 3B**). However, treatment of macrophages with c-di-AMP alone did not induce IFN-β production and *Ifnb1*, *Cxcl10*, and *Ifit1* expression (**Fig. 3B**). These results observed from RAW264.7 macrophages were confirmed by using BMDMs (**Fig. 3C**) and human macrophages (THP-1) (**Fig. 3D**).

**Fig. 3.**
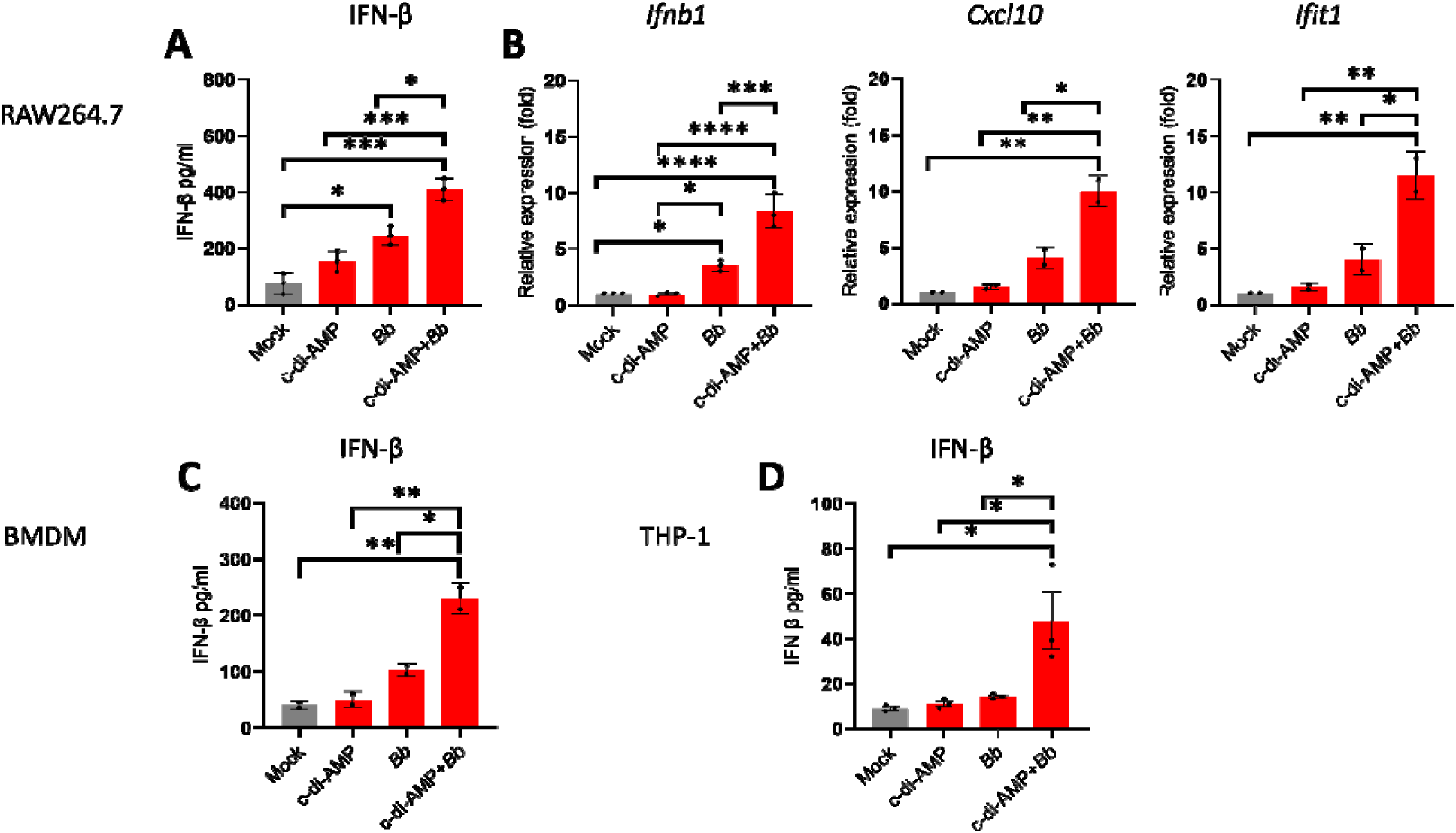
Exogenous c-di-AMP enhances *B. burgdorferi-*induced IFN-I response *in vitro* and *ex vivo.* RAW264.7, BMDMs and human THP-1 cells were infected with wild-type (WT) *B. burgdorferi* 5A4NP1 in the absence and presence of exogenous c-di-AMP (1µg/ml) at an MOI of 10. After 24 hours, supernatants and macrophage cells were collected. The IFN-β protein level in the supernatant of infected RAW264.7 **(A),** BMDMs (**C**), and THP-1 cells (**D**) were measured by ELISA. The transcript levels of *Ifnb1*, *Cxcl10,* or *Ifit1* (**B**) in infected RAW264.7 cells were determined by qRT-PCR analysis (n=3) and normalized with the housekeeping gene *actin*. The levels of gene expression relative to mock-infected control were reported. The data represent the mean ± SD of at least three experiments (n=3). Statistical significance was assessed using the one-way ANOVA, with \**p* < 0.05, \*\**p* < 0.01 and \*\*\**p* < 0.001.

### Exogenous c-di-AMP enhances *B. burgdorferi-*induced IFN-I response during mouse infection

We next sought to investigate the role of *B. burgdorferi* c-di-AMP in IFN-I production. during infection. Ideally, the *cdaA* mutant would be used for this purpose; however, the *cdaA* mutant was unable to infect mice. As an alternative approach, groups of C3H/HeN mice were subcutaneously infected with *B. burgdorferi* either in the presence of c-di-AMP (25 µg/mouse) or PBS (control). After 1 and 7 days of infection, mouse skin tissue at the site of inoculation was harvested and the mRNA levels of *Cxcl10* and *Ifit1* were determined using Quantitative real-time PCR (qRT-PCR) (**Fig. 4A**). After one day of infection, neither c-di-AMP treatment nor *B. burgdorferi* infection alone showed significant induction of *Cxcl10* and *Ifit1* expression (**Fig. 4B**). However, when *B. burgdorferi* co-inoculated with exogenous c-di-AMP, the *Cxcl10* and *Ifit1* expressions were significantly increased (**Fig. 4B**). At day 7 of infection, *B. burgdorferi* infection showed *Cxcl10* and *Ifit1* expression, and exogenous c-di-AMP further enhanced *Cxcl10* and *Ifit1* expression (**Fig. 4C**). This result suggests that during mammalian infection, c-di-AMP plays an important role in *B. burgdorferi*-induced IFN-I response.

**Fig. 4.**
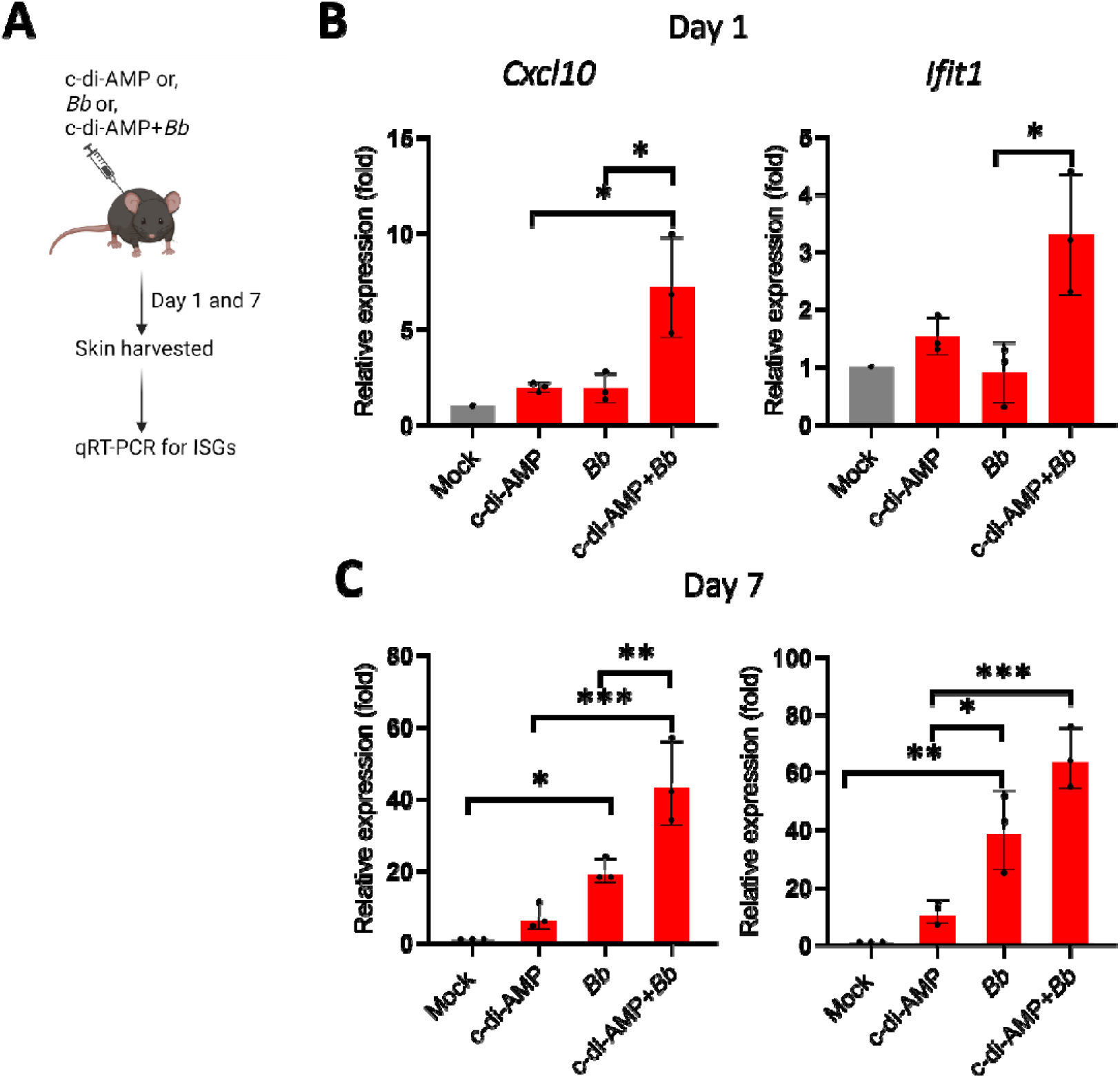
Exogenous c-di-AMP enhances *B. burgdorferi*-induced IFN-I response in mice. **(A).** A diagram depicting the experimental procedure. Groups of mice were infected with wild-type *B. burgdorferi* (10^5 /mouse, subcutaneous) either in the absence or presence of exogenous c-di-AMP (25µg/mouse). 24 hours after infection, mice were sacrificed, and skin tissues were isolated at the site of infection. **(B)** qRT-PCR analysis of ISG genes (*Cxcl10* and *Ifit1*) expression in isolated mouse skin tissues was performed on days 1 **(B)** and 7 **(C).** Data were normalized with the housekeeping gene *actin*. The levels of gene expression relative to mock-infected control were reported. The data are mean ± SD of three mice in each group (n=3/group). Statistical significance was assessed using the one-way ANOVA, with \**p* < 0.05, \*\**p* < 0.01 and \*\*\**p* < 0.001.

### *B. burgdorferi* activates STING signaling pathway

*B. burgdorferi* induces IFN-I response independent of the TLR2 signaling pathway. To determine the role of the STING signaling pathway in *B. burgdorferi*-induced IFN-I response, we first examined the phosphorylation of STING, TBK1, and IRF3. The immunofluorescence assay-results showed that *B. burgdorferi* infection-induced phosphorylation of STING, TBK1, and IRF3 (**Fig. 5A**). Similarly, Western blot analyses revealed a dose-dependent increase in the phosphorylation of STING, TBK1, and IRF3 in macrophages, correlating with higher multiplicity of infection (MOI) of *B. burgdorferi* (**Fig. 5B**). We then examined the impact of STING KO on IFN-I induction. The results showed that STING KO significantly reduced the IFN-β protein level and the *Ifnb1* RNA levels (**Fig. 6A**, left panel). As an alternative approach, the effects of inhibitors H-151 against STING, or BX795 against TBK1, were examined. Treatment with H-151 or BX795 dramatically reduced both mRNA and protein levels of IFN-β induced by *B. burgdorferi* infection (**Fig. 6A**, middle and right panel). In contrast, Poly I: C, which induces IFN-I response by TLR-dependent pathways but independent of STING, showed no effect on the IFN-β protein level and the *Ifnb1* RNA levels in RAW and STINGKO macrophages as well as in the presence of STING and TBK1 inhibitors (**Fig. 6B**). These results are consistent with the recent finding that *B. burgdorferi*-induced IFN-I response is STING-dependent (Farris *et al*., 2023).

**Fig. 5.**
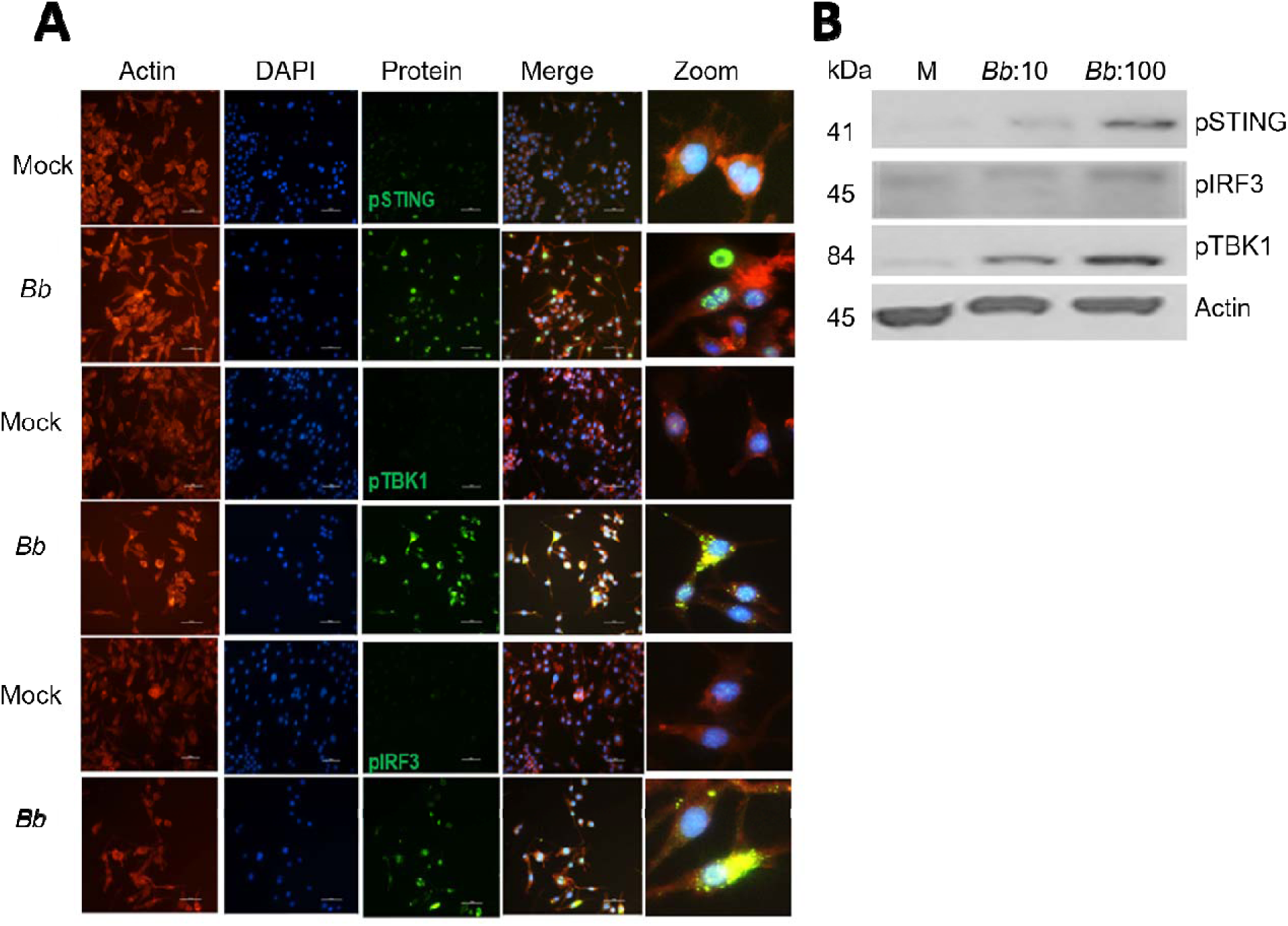
*B. burgdorferi* infection to macrophages resulted in the phosphorylation of STING, TBK1, and IRF3. **(A)** Fluorescence microscopy merged (Blue+Green+ Red channels) images of macrophages that were fixed after 24 hpi with *B. burgdorferi* (MOI of 10) and stained with anti-pTBK1, anti-pSTING, anti-pIRF3 antibodies. Red, Actin (AF653); Blue, Nuclei (DAPI); Green, pTBK1, pSTING, or pIRF3 (AF488). **(B)** Western blot analysis of pSTING, pTBK1, pIRF3, and Actin (loading control) of RAW264.7 cells infected with wild-type *B. burgdorferi* for 24 hours at an MOI of 10 or 100. The data are representative images of two independent experiments.

**Fig. 6.**
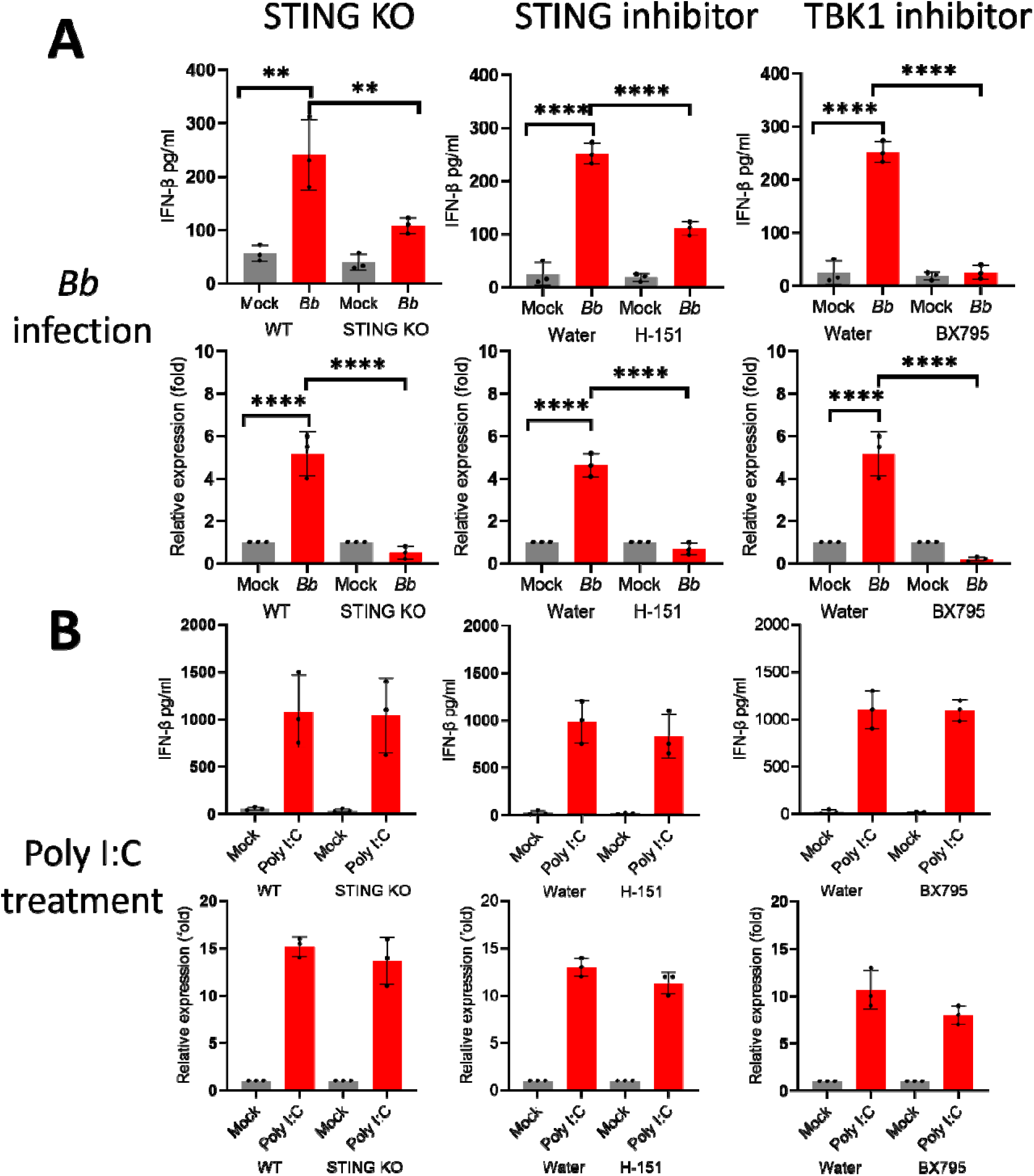
*B. burgdorferi-*induced IFN-β is dependent on the STING signaling pathway. RAW264.7 or STING KO macrophages treated with STING inhibitor H-151 (100 ng/ml) or TBK1 inhibitor BX795 (100 ng/ml), were infected with wild-type (WT) *B. burgdorferi* 5A4NP1 at an MOI of 10 (**A**). As controls, cells were treated with poly I: C (**B**). After 24 hours, supernatants and macrophage were collected. The IFN-β protein levels were measured in supernatants (**A** and **B,** top panel) and *Ifnb1* RNA levels were measured in macrophages and represented as relative to mock-treated (**A** and **B,** lower panel). The data represent the mean ± SD of at least three experiments (n = 3). Statistical significance was assessed using one-way ANOVA, with \*\**p* < 0.01, \*\*\**p* < 0.001, and \*\*\*\**p* < 0.0001.

### c-di-AMP plays a crucial role in STING activation by *B. burgdorferi*

To determine the role of c-di-AMP in the activation of STING signaling by *B. burgdorferi*, macrophages were infected with wild-type *B. burgdorferi*, the Δ*cdaA* mutant, or the Δ*dhhP* mutant spirochetes. The number of pSTING-positive macrophages was quantified using immunofluorescence assays. The results showed that a higher percentage of pSTING-positive macrophages was observed with the Δ*dhhP* infection and a lower percentage with the Δ*cdaA* infection (**Fig. 7A and 7B**). To determine whether IFN-I induction by the supernatants of *B. burgdorferi* culture is dependent on STING, SVPD-treated, or untreated *B. burgdorferi* supernatants were incubated with RAW or STING KO cells for 24 hrs. The result shows that when exposed to the supernatants of *B. burgdorferi* culture, STING KO cells produced lower levels of IFN-β (**Fig. 7C**) and IRF (interferon response factor) (**Fig. 7D**) than that of RAW cells, suggesting that induction of IFN-I response by the supernatant of *B. burgdorferi* culture is STING dependent. Interestingly, levels of IFN-β and IRF were further reduced when STING KO cells were exposed to SVPD-treated *B. burgdorferi* supernatants (**Fig. 7C. 7D**), suggesting that c-di-AMP also triggers a STING-independent induction of the IFN-I response.

**Fig. 7.**
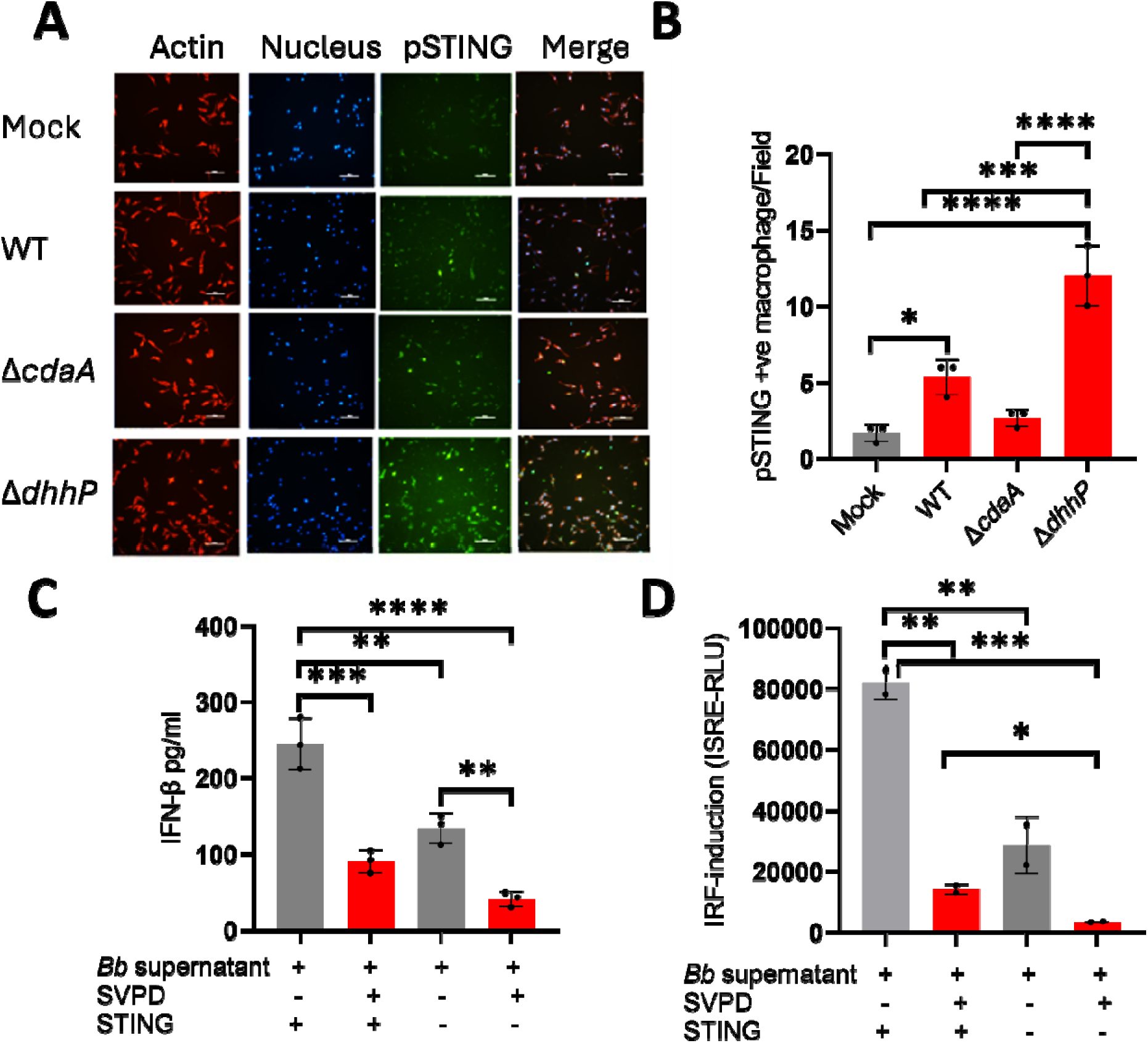
STING activation by *B. burgdorferi* is dependent on c-di-AMP. (**A**) Fluorescence microscopy merged (Blue+Green+ Red channels) images of macrophages stained with anti-pSTING antibodies. RAW 264.7 macrophages were infected with wild-type (WT) *B. burgdorferi*, Δ*cdaA*, or Δ*dhhP* strains and were fixed after 24 hpi. Nuclei-Blue (DAPI), pSTING-Green (AF488), Actin-Red (AF653). (**B**) Quantitative analysis of pSTING-positive macrophage. (**C**) Levels of IFN-β were determined by ELISA in culture media in WT or STING KO macrophage (at 24 hpi) incubated with *B. burgdorferi* culture supernatants pre-treated with or without phosphodiesterase (SVPD). (**D**) IRF induction was measured by luciferase reporter assay in macrophage supernatants treated with the *B. burgdorferi* culture supernatants (with or without SVPD treatment). The data represent the mean ± SD of at least three experiments (n = 3). Statistical significance was assessed using the one-way ANOVA, with \**p* < 0.05, \*\**p* < 0.01, \*\*\**p* < 0.001, and \*\*\*\**p* < 0.0001.

### c-di-AMP is a key contributor to *B. burgdorferi*-induced interferon-stimulated gene expression

*B. burgdorferi* infection induces host ISG expression, which enhances the pathogenicity and plays a crucial role in the development of Lyme arthritis (Crandall *et al*., 2006, Miller *et al*., 2010, Miller *et al*., 2008). To determine the role of c-di-AMP of *B. burgdorferi* in ISGs expression, RAW or STING KO macrophages were infected with wild-type *B. burgdorferi* or Δ*cdaA* strains, and ISGs expressions were determined 24 hours post-infection. The result showed that *B. burgdorferi* infection-induced expressions of all six ISG tested in RAW macrophages and STING KO virtually abolished these expressions (**Fig. 8A** and **8B**), indicating that ISG induction was STING dependent. Furthermore, the Δ*cdaA* mutant strain induced a significantly lower level of ISG expression than that of wild-type *B. burgdorferi*, suggesting that c-di-AMP contributes significantly to *B. burgdorferi*-induced ISG expression (**Fig. 8C**).

**Fig. 8.**
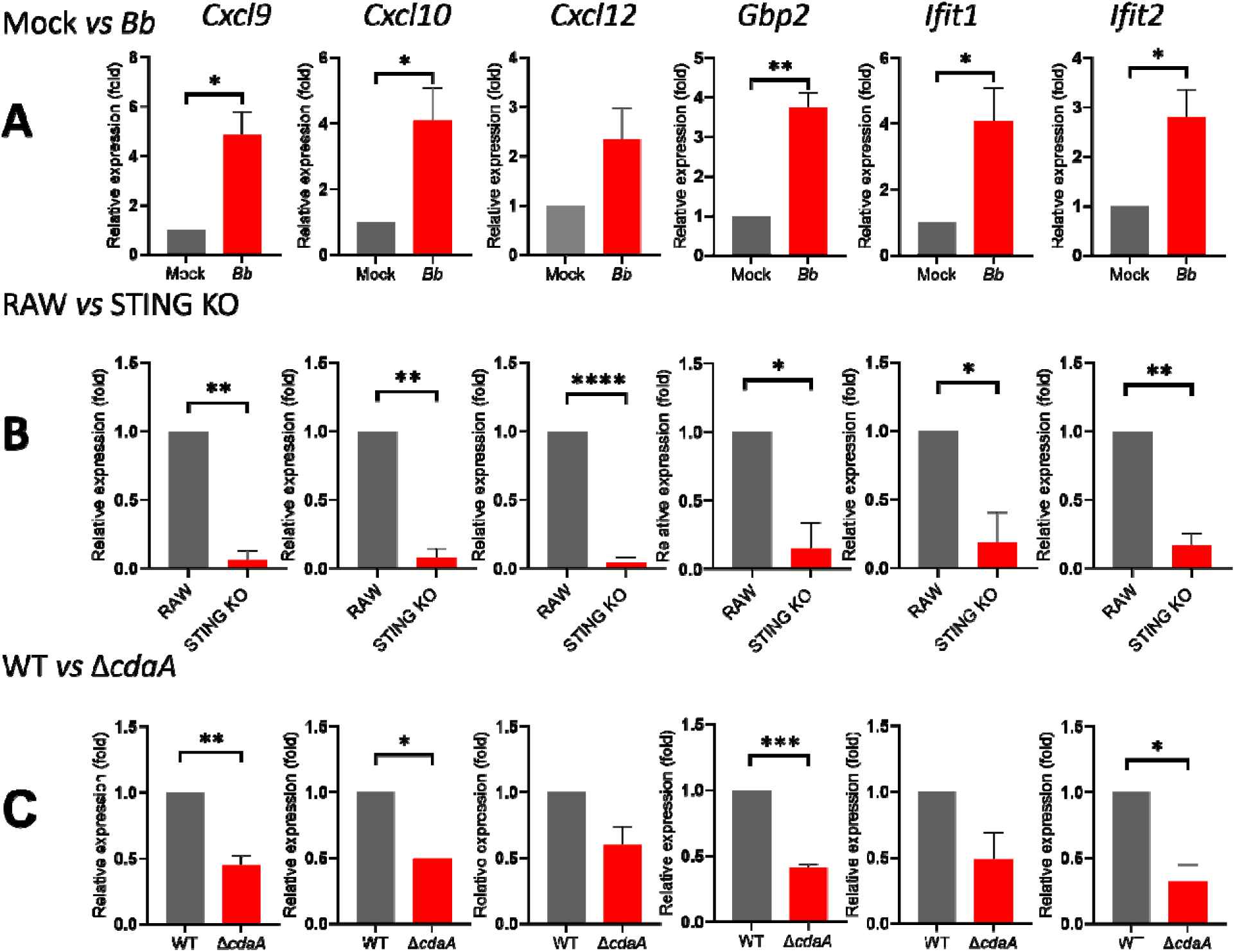
The induction of interferon-stimulated genes (ISG) in macrophages by *B. burgdorferi* is dependent on both the level of c-di-AMP and the STING signaling. (A) qRT-PCR analysis of gene expression in macrophage uninfected (mock) or infected with *B. burgdorferi* for 24 hours at an MOI of 10 **(B)** qRT-PCR analysis of gene expression in WT and STING KO cell lines infected with *B. burgdorferi* for 24 hours at an MOI of 10. **(C)** qRT-PCR analysis of gene expression in WT cell line infected with WT or mutant strain of *B. burgdorferi* for 24 hours at an MOI of 10. After infection, cells were collected, and total RNA was isolated. qRT-PCR was performed using primers for CXCL9, CXCL10, CXCL12, GBP2, IFIT1 and IFIT3. Actin was used as a housekeeping gene. The data represent the mean ± SD of at least three experiments (n = 3). Statistical significance was assessed using the two-tailed Student’s t-test, with \**p* < 0.05, \*\**p* < 0.01, \*\*\**p* < 0.001, and \*\*\*\**p* < 0.0001.

**Fig. 9.**
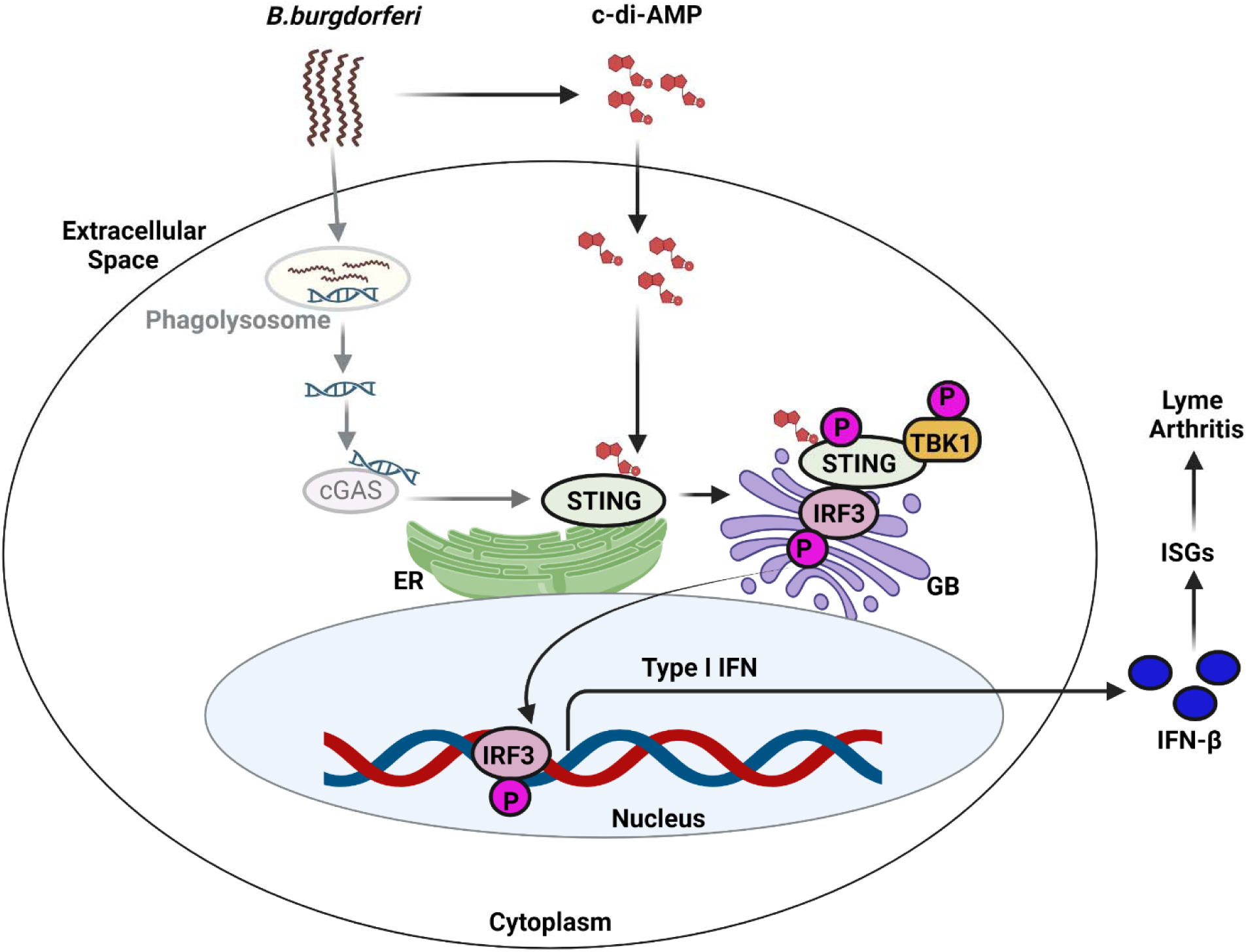
A proposed model of the mechanism of induction of IFN-I response by *B. burgdorferi* in macrophage. The second messenger c-di-AMP secreted by *B. burgdorferi* binds directly to the STING protein, resulting in the activation of downstream proteins and induction of IFN-I response. In addition, *B. burgdorferi* DNA released from the phagolysosome may interact with cGAS and activate STING. ER, endoplasmic reticulum; GB, Golgi bodies; cGAS, cyclic GMP-AMP synthase; IRF3, interferon response factor 3; TBK1, TANK-binding kinase; ISGs, interferon stimulator genes.

## DISCUSSION

*B. burgdorferi* induces a robust IFN-I response that plays a key role in the development of Lyme arthritis (Miller *et al*., 2008, Miller *et al*., 2010). Despite this, the specific *B. burgdorferi*-derived PAMPs that trigger IFN-I responses remain largely undefined (Oosting *et al*., 2010, Wooten *et al*., 2002). Previous studies have suggested that *B. burgdorferi* extracellular components secreted to the culture supernatant elicit strong IFN-I response (Miller *et al*., 2010). However, the identity of such PAMP remains unknown. In this study, we uncovered that *B. burgdorferi* secretes c-di-AMP that serves as a key PAMP to induce host IFN-I response via activating the STING signaling pathway. Several lines of evidence support the conclusion that c-di-AMP is a secreted PAMP that induces IFN-I response by *B. burgdorferi*. Firstly, c-di-AMP was readily detected in the supernatant of *B. burgdorferi* culture (**Fig. 1B**). Secondly, the Δ*cdaA* and Δ*dhhP* mutant induced lowed and elevated levels of IFN-I response, respectively (**Fig. 2**). Thirdly, the levels of c-di-AMP in culture supernatants from various *B. burgdorferi* strains correlated with the levels of IFN-I induction, and treatment of the supernatants with the exogenous phosphodiesterase SVPD resulted in a reduction in IFN-I production (**Fig. 2**). Fourthly, addition of exogenous c-di-AMP to the co-culture of *B. burgdorferi* and macrophages enhances IFN-I response (**Fig. 3**). Finally, co-inoculation of c-di-AMP and *B. burgdorferi* enhances IFN-I response at the site of inoculation in mice (**Fig. 4**). Collectively, our data provide strong evidence that c-di-AMP is a secreted PAMP produced by *B. burgdorferi*, triggering IFN-I response.

*B. burgdorferi* induces IFN-I response independent of MyD88 and TRIF signaling pathway (Miller *et al*., 2008). Recently, Farris *et al*., demonstrated that *B. burgdorferi* induces an IFN-I response by activating the cGAS-STING signaling pathway (Farris *et al*., 2023). Consistent with that study, our present study also showed the induction of IFN-I response by *B. burgdorferi* is STING-dependent (**Fig. 6**). Furthermore, we identified that c-di-AMP is one of the key PAMPs of *B. burgdorferi* that activates the STING signaling pathway. We provided several pieces of evidence to support this finding. Firstly, a decrease in macrophage positive for activated STING was observed on infection with Δ*cdaA* strain while an increase was observed on infection with Δ*dhhP* strain as compared to the wild-type strain (**Fig. 7A and 7B**). Secondly, a marked reduction in IFN-I response was observed when *B. burgdorferi* supernatant treated with SVPD was used to expose the wild-type macrophages, and the level of IFN-I response was further reduced in the STING KO macrophages (**Fig 7C and 7D**). Thirdly, a decrease in the level of ISG expression was observed in the STING KO macrophages as compared to wild-type macrophages infected with *B. burgdorferi* (**Fig. 8**).

The discovery that c-di-AMP triggers the host type I interferon (IFN-I) response was first made in the intracellular bacterium *Listeria monocytogenes* by Woodward *et al*. (Woodward *et al*., 2010). It is now well-established that bacterial cyclic dinucleotides (c-di-AMP, c-di-GMP, 3’3’-cGAMP), subsequently, the host 2’3’-cGAMP, activate STING and induce IFN-I production (Liu *et al*., 2022, Zaver & Woodward, 2020, Wright & Bai, 2023, Stülke & Krüger, 2020, Gries *et al*., 2016, Andrade *et al*., 2016, Barker *et al*., 2013, Woodward *et al*., 2010, Burdette *et al*., 2011, Moretti *et al*., 2017). Notably, previous studies have primarily focused on the function of c-di-AMP in Gram-positive bacteria, with the exception of *Chlamydia trachomatis* (*C. trachomatis*), an obligate intracellular Gram-negative bacterium. *C. trachomatis* has been shown to produce c-di-AMP, which is crucial for STING activation and the IFN-I response (Barker *et al*., 2013). However, that study did not use a c-di-AMP-deficient *C. trachomatis* mutant to fully satisfy molecular Koch’s postulates. As *B. burgdorferi* is an extracellular, Gram-negative-like spirochete, the findings in the current study broaden our understanding of c-di-AMP’s role across the bacterial kingdom.

One interesting observation in this study is that c-di-AMP treatment alone induces no or low IFN-I response, suggesting that the effect of c-di-AMP requires the presence of *B. burgdorferi* (or additional components found in *B. burgdorferi* supernatant) (**Fig. 3 and 4**). Unlike intracellular pathogens, *B. burgdorferi*-secreted c-di-AMP must enter host cells to bind and activate STING. Indeed, previous reports on other extracellular bacteria such as Group B *Streptococcus*, observed that the effect of c-di-AMP on IFN-β reduction was only observed when transfection was performed (Andrade *et al*., 2016). The observation that *B. burgdorferi* promotes the effect of c-di-AMP on STING activation is intriguing. One possibility is that *B. burgdorferi* itself or its component creates a permissive environment for c-di-AMP uptake by host cells. It has reported that mediators released by *B. burgdorferi* can directly enhance mammalian cell permeability (Zhou *et al*., 2008). It is perceivable that increased host cell permeability facilitates the intracellular passage of c-di-AMP and enhances its immunostimulatory effects. *B. burgdorferi* may also affect host cell transporters SLC19A1 and SLC46A2, which are capable of transporting cyclic-di-nucleotides to the cytosol, leading to an increased uptake of c-di-AMP (Luteijn *et al*., 2019, Ritchie *et al*., 2019, Cordova *et al*., 2021). Another possibility is that *B. burgdorferi* or its components may inhibit the cell receptor ENPP1, which has been reported capable of degrading cyclic-di-nucleotides (Li *et al*., 2014, Onyedibe *et al*., 2019), resulting in higher c-di-AMP levels in the environment. Other possible mechanisms, such as pinocytosis or endocytosis, may be manipulated by *B. burgdorferi* and contribute to an increased uptake of c-di-AMP by host cells (Liu *et al*., 2019). These findings collectively suggest that although c-di-AMP is a negatively charged polar compound, it can efficiently cross host cell membranes in the presence of *B. burgdorferi.* This enhanced uptake and subsequent immune activation highlight a potential strategy employed by *B. burgdorferi* to manipulate host immune responses. Further study into these mechanisms will provide deeper insights into how *B. burgdorferi* facilitates the intracellular uptake of c-di-AMP and enhances the host’s IFN-I response, potentially revealing novel targets for therapeutic intervention.

Another interesting observation in this study is that c-di-AMP appears to trigger a STING-independent activation in addition to STING-dependent activation (**Fig. 7C** and **7D**). These results suggest that there might be alternative sensors or signaling molecules in the host cells that respond to c-di-AMP, leading to the induction of the IFN-I response. Although receptors such as cGAS and IFI16 are known to activate the IFN-I response in a c-di-AMP-independent and STING-dependent manner, the existence of signaling pathways that are dependent on c-di-AMP but independent of the STING protein remains unexplored (Unterholzner *et al*., 2010). Our findings highlight the need for future studies to investigate the molecular mechanisms underlying this STING-independent activation. Identifying these components could provide deeper insights into the innate immune responses elicited by *B. burgdorferi* and possibly other pathogens.

Previous studies have shown that IFN-I induction in response to several bacterial pathogens including *Mycobacterium tuberculosis*, *L. monocytogenes*, *C. trachomatis*, and Group B Streptococcus is primarily driven by the host recognition of bacterial DNA, whereas bacterial c-di-nucleotides play a minor role in stimulating the IFN-I response (Hansen *et al*., 2014, Collins *et al*., 2015, Wassermann *et al*., 2015, Watson *et al*., 2015, Andrade *et al*., 2016, Zhang *et al*., 2014). However, in the case of *B. burgdorferi*, c-di-AMP appears to play a significant role, as the *cdaA* mutant exhibited over 50% reduction of IFN-β reduction (**Fig. 1A**). On the other hand, other PAMPs of *B. burgdorferi* also contribute to the induction of the IFN-I response, as deletion of *cdaA* did not completely abolish IFN-β production (**Fig. 1A**). Indeed, Farris *et al*, demonstrated the co-localization of *B. burgdorferi* and cGAS (Farris *et al*., 2023). In addition, *B. burgdorferi* infection may result in host microbial or nuclear DNA releasing into the cytosol and activate cGAS-STING axis, as reported in other bacterial infections (Watson *et al*., 2015, West & Shadel, 2017, Zhou *et al*., 2021). Furthermore, *B. burgdorferi* produces another cyclic dinucleotide, c-di-GMP. Studies have shown that c-di-GMP (He *et al*., 2011, Kostick *et al*., 2011), which may also contribute STING activation and IFN-I production.

One limitation of this study is the inability to establish a direct correlation between c-di-AMP levels and the IFN-I response in an *in vivo* setting. Although we successfully generated mutant strains of *B. burgdorferi* producing higher and lower levels of c-di-AMP, both elevated levels and lack of c-di-AMP proved deleterious to *B. burgdorferi* under *in vivo* conditions. This suggests that a moderate level of c-di-AMP is essential for the establishment and maintenance of infection by *B. burgdorferi* in its host. The development of conditional mutants could be a viable approach in this scenario. Conditional mutants would allow us to modulate c-di-AMP levels in a controlled manner during infection, thereby providing insights into the interplay between c-di-AMP and the host immune response. Such an approach would enable us to overcome the limitations posed by the deleterious effects of extreme c-di-AMP levels, thereby facilitating a better understanding of its role in pathogenesis. Despite these challenges, our observations indicate that the exogenous addition of c-di-AMP leads to an increased induction of the IFN-I response in both *in vivo, ex vivo, and in vitro* conditions. This suggests that c-di-AMP levels indeed modulate the innate immune response, emphasizing its significance as a critical PAMP in the host-pathogen interaction in the context of *B. burgdorferi* pathogenesis.

In summary, our findings underscore the critical role of the *B. burgdorferi*-secreted c-di-AMP as a key PAMP in inducing the IFN-I response via the STING signaling pathway. Our finding showing that macrophages exposed to heat-killed *B. burgdorferi* has a significant reduction in IFN-β production (**Fig. 2C and 2F**), support the notion that unlike classical PAMP, c-di-AMP is a vita-PAMP (Moretti *et al*., 2017), i.e., a signature of live *B. burgdorferi*. Noted that previous studies also observed that human monocytes exhibit a more robust IFN-I response when infected with live *B. burgdorferi* compared to exposure to bacterial lysates (Salazar *et al*., 2009) or heat-killed spirochetes (Petzke *et al*., 2009). Unlike classical PAMPs triggering an immediate immune response, vita-PAMPs associated with viable pathogens, offer the added benefit of distinguishing live pathogens from dead ones, leading to a more targeted and efficient immune response while reducing unnecessary inflammation. This dual role can be beneficial to the host but may also allow bacteria to evade immune detection by modulating immune responses for persistent infection. Future studies will elucidate the roles of c-di-AMP and IFN-I production in *B. burgdorferi* pathogenesis and investigate the therapeutic efficacy of targeting c-di-AMP production as a novel strategy for the prevention/treatment of Lyme disease.

## MATERIALS AND METHODS

### Ethics Statement

All experiments were approved by the Indiana University School of Medicine’s Institutional Animal Care and Use Committee and performed following the institutional guidelines for animal use. Mice were housed under specific-pathogen-free conditions at LARC (Laboratory Animal Research Centre). All efforts were made to reduce animal suffering during this study.

### Cells and culture conditions

Murine Raw 264.7-derived macrophages, RAW-Lucia ISG cells, were provided by Dr. Hernim Sintim at Purdue University, Indianapolis, Indiana. RAW-Lucia ISG-KO STING (STING KO) was procured from Invivogen. The L929 cell line was provided by Dr. Randy Brutkiewicz, Indiana University, Indianapolis, Indiana. The human THP1 cell line was used from our lab cell lines inventory. Raw 264.7-derived macrophages and L929 cell lines were cultured in DMEM supplemented with 10% heat-inactivated FBS,100 U/ml penicillin, and 100 μg/ml streptomycin. The human THP-1 cell line was cultured in RPMI 1640 supplemented with 10% heat-inactivated FBS, 100Uml penicillin, 100 μg/ml streptomycin, and 25 mM HEPES. For the infection study, the THP-1 cell line was first differentiated using RPMI 1640 complete media + Phorbol 12-myristate-13-acetate (PMA, 5ng/ml) for 24 hours and then allowed to rest in RPMI 1640 complete media for another 24 hours before infection.

Primary murine bone marrow-derived macrophages (BMDMs) were prepared from 4-6-week-old C3H/HeN mice, following a modified protocol described previously (Toda *et al*., 2021). Briefly, bone marrow was isolated from the femur and tibiae in DMEM supplemented with 10% FBS and flushed through a syringe with a 20-gauge needle. The cells were washed once in PBS and then seeded in petri dishes containing 10 ml of DMEM supplemented with 10% FBS, 2 mM L-glutamine, 2% HEPES, 1 mM sodium pyruvate, 100 U/ml penicillin,100 μg/ml streptomycin, and 30% L929-conditioned media. The cultures were maintained at 37°C with 5% CO2. After 4 days of incubation, fresh medium was added. On day 6, macrophages were detached from the plate surface, and cells were assessed for purity by double staining with anti-CD11b (BioLegand, Clone: M1/70, catalog, 101212) and anti-F4/80 (BioLegend, Clone: BM8, catalog 123109) macrophage markers using flow cytometric analysis before being utilized for infection studies.

### Mice

C3H/HeN mice were purchased from Jackson Laboratory (Bar Harbor, ME) and maintained at the Indiana University, LARC facility. Both male and female mice aged 4-6 weeks were used for all the experiments. Mice were housed under specific-pathogen-free conditions at LARC (Laboratory Animal Research Centre). All experiments were approved by the Indiana University School of Medicine’s Institutional Animal Care and Use Committee and performed following the institutional guidelines for animal use.

### Reagents and antibodies

Antibodies were purchased as follows: anti-pIRF3 (Cell signaling, 4947S), anti-pTBK1 (Cell signaling, 5483S), anti-pSTING (Cell Signaling, 72971S), Rabbit IgG Isotype control (Cell signaling), anti-mouse Alexa fluor-488 (Cell Signaling), anti-mouse HRP conjugate (Cell signaling), Alexa Fluor 594 anti-Rabbit IgG (Cell Signaling, 8889S), Alexa Fluor 488 anti-Rabbit IgG (Cell Signaling, 4412S), PE anti-mouse F4/80 (BioLegend, Clone: BM8, 123109), APC anti-mouse/human CD11b (BioLegand, Clone: M1/70, 101212). The following reagents were used: PMA (Sigma-Aldrich, P1585) SVPD (Millipore Sigma, P3243), c-di-AMP ELISA kit (Cayman Chemical, 501960), c-di-AMP (InvivoGen, tlrl-nacda), Poly I: C (InvivoGen, tlrl-picw), H-151 (InvivoGen, inh-h151), BX795 (InvivoGen, tlrl-bx7-2)

### *B. burgdorferi* strains and culture conditions

The *B. burgdorferi* strains utilized in this study comprised the wild-type strain 5A4NP1, obtained from our laboratory collection. The mutant strain lacking the gene for the synthesis of phosphodiesterase (Δ*dhhP*) was previously generated in our laboratory (Ye *et al*., 2014). The mutant strain lacking the gene for the synthesis of c-di-AMP (Δ*cdaA*) was constructed using the allele exchange method (Zhang *et al*., 2020). Briefly, the suicide vector pMP04 was first constructed, which contains a 4 kb DNA fragment of *cdaA* region, except that the cdaA gene was replaced by an *aacC1* marker (which confers gentamycin-resistant) (**Fig. S1A**). pMP04 was then transformed into 5A4NP1, and gentamycin-resistant transformants were analyzed by PCR to confirm the correct *cdaA* deletion (**Fig. S1B**). For complementation (the mutant replacement), the suicide vector pMP24 was constructed, which contains 4 kb DNA fragment of wild-type *cdaA* region, except that an *aadA* marker (which confers streptomycin-resistant). Streptomycin-resistant and gentamycin-sensitive transformants were analyzed by PCR to confirm the correct *cdaA* restoration (**Fig. S1B**).

All strains were cultured in the standard Barbour-Stoenner-Kelly II (BSK-II) medium with appropriate antibiotics at 37°C and pH 7.5 in a 5% CO2 incubator until the late log phase. *B. burgdorferi* titer was determined by counting spirochetes under a dark-field microscope (Olympus America Inc., Center Valley, PA). Antibiotics were employed at the following concentrations: streptomycin and gentamycin at 50 µg/ml, and kanamycin at 100 µg/ml. Heat-killed *B. burgdorferi* were generated by immersing the culture in a 65°C water bath for 40 minutes as described previously (Petzke *et al*., 2009). The efficacy of heat-killing was confirmed by culturing the spirochetes in BSK-II media at 37°C for 1 week.

### Quantification of c-di-AMP level by ELISA

Wild-type and mutant strains of *B. burgdorferi* were cultured in BSK-II media at 37°C until reaching the late log phase. The cultures were centrifuged at 8,000 rpm (revolutions per minute) for 10 minutes, and the supernatants and pellets were collected separately. The pellets were washed three times with PBS and subsequently lysed in B-PER reagent (Thermo Scientific, catalog 78243). Protein concentration was determined using the Bardford Assay Reagent (Thermo Scientific, catalog 1863028).

To detect extracellular c-di-AMP levels, the collected supernatants were filtered through a 0.22 µm filter and diluted 5-fold in DMEM. In certain experiments, supernatants were treated with either water or SVPD (Millipore Sigma, catalog P3243) for 2 hours at RT (room temperature) (2 IU/ml of culture), as previously described (Barker *et al*., 2013). The levels of c-di-AMP in the lysates and supernatants were quantified using a c-di-AMP ELISA kit (Cayman Chemical, catalog 501960), following the manufacturer’s instructions.

### *B. burgdorferi* culture supernatant preparation for macrophage infection

Culture supernatants from *B. burgdorferi* were prepared following a previously described protocol (Miller *et al*., 2010). Briefly, wild-type and mutant strains of *B. burgdorferi* (Δ*cdaA* and Δ*dhhP*) were cultured in BSK-II media until reaching the late log phase. The supernatants were collected by centrifugation at 8,000 rpm for 10 minutes, followed by filtration through a 0.22 µm filter (Fisher Scientific, catalog 09-719C). The filtered supernatants were then diluted 5-fold in DMEM and used to infect macrophages. As a control, BSK-II media without *B. burgdorferi* was similarly diluted in DMEM and used to treat the macrophages.

### Infection of macrophages with *B. burgdorferi*

RAW-Lucia ISG (InvivoGen, catalog rawl-isg) and RAW-Lucia ISG-KO-STING cells (InvivoGen, catalog rawl-kostg) were derived from the murine RAW 264.7 macrophage cell line through stable integration of an interferon regulatory factor (IRF)-inducible secreted luciferase construct. These cells, in conjunction with bone marrow-derived macrophages (BMDMs) isolated from C3H/HeN mice, were infected with wild-type (live or heat-killed) or mutant strains of *B. burgdorferi* at a MOI of 10 for 3 and 24 hours. In certain experiments, spent culture supernatants from wild-type and mutant strains of *B. burgdorferi* were treated with SVPD for 2 hours at RT and subsequently used to expose the macrophages. Additionally, the macrophages were treated with either water (mock), STING inhibitor, H-151 (1 µg/ml, InvivoGen, catalog inh-h151), TBK1 inhibitor, BX795 (100 ng/ml, InvivoGen, catalog tlrl-bx7-2) for 1 hour and then infected with either *B. burgdorferi* or treated with Poly I: C (100 ng/ml) (InvivoGen, tlrl-picw), for 24 hours. To check the effect of c-di-AMP, macrophages were infected with *B. burgdorferi* at MOI 10 or mock-treated in the presence of c-di-AMP (1 µg/ml) (InvivoGen, catalog tlrl-nacda) or water for 24 hours. Following infection, macrophages supernatants were collected, centrifuged, and utilized for the measurement of interferon-beta (IFN-β) by enzyme-linked immunosorbent assay (ELISA). The cells were collected for the RNA extraction as described below.

### IFN-β detection by ELISA

ELISA for IFN-β was conducted with macrophage supernatants using a cytokine ELISA kit specific for mice following the manufacturer’s instructions. In brief, ELISA plates were coated with purified antibody (3 µg/ml) overnight at 4°C (BioLegend, catalog 519202). The next day, plates were washed thrice with PBS-T and blocked with 2% BSA for 2 hours at RT. After washing, plates were incubated with samples (100 µl) overnight at 4°C, followed by washing. Subsequently, plates were incubated with biotin-conjugated secondary antibody for 1 hour at RT (BioLegend, catalog 508105). The plates were washed with PBS-T and incubated with Avidin-peroxidase for 30 minutes at RT (Sigma-Aldrich, catalog A3151). After washing with PBST, TMB substrate (100 µl, BioLegend, catalog 421101) was added to each well, followed by the addition of stop solution (100 µl). The plates were read at an absorbance of 450 nm. The cytokine concentration was determined by a standard curve generated with a known concentration of IFN-β (BioLegand, catalog 581309).

### Immunofluorescence staining

RAW cells were seeded at 10^5 cells/well in a 24-well cell culture dish containing glass coverslips and infected with *B. burgdorferi* at an MOI of 10. At 24 hours post-infection (hpi), infected and mock-treated cells were washed with PBS three times and fixed with 4% paraformaldehyde (PFA) for 10 minutes at RT. After three washes with PBS, cells were permeabilized with 0.1% Triton X-100 for 10 minutes at RT. Subsequently, cells were washed with PBS and blocked with 2% BSA for 1 hour at RT. The cells were incubated with primary antibodies, anti-pSTING (Cell Signaling, catalog 62912S), pTBK1 (Cell signaling, catalog 5483S), or pIRF3 (Cell Signaling, catalog 29047S) antibodies (1:100 dilution) overnight at 4°C in blocking buffer. Following this, cells were washed with PBS and incubated with an anti-Rabbit IgG Alexa Fluor 488 (Cell Signaling, catalog 4412S) /594 (Cell Signaling, catalog 8889S)-conjugated secondary antibody (1:1000) for 1 hour at RT, followed by another round of washing with PBS. Finally, cells were mounted on glass slides with a Vectashield anti-fade mounting medium containing DAPI (Vector Laboratories, Burlingame, CA). The slides were imaged using a Nikon Scanning microscope with 20X and 60X objectives in the green, blue, and red channels.

### Western blotting analysis

For western blot analysis, macrophages were infected with *B. burgdorferi* at an MOI of 10 or 100. After 24 hpi, the infected cells were washed twice with PBS and lysed in cell lysis buffer (Cell Signaling, catalog 9803) containing protease inhibitor (Cell Signaling, catalog 5871S) and phosphatase inhibitor (Cell Signaling, catalog 5870S). The cells were fully lysed by pipetting followed by sonication. Protein quantification was performed, and samples were resuspended in 3X Laemmli buffer and heated for 5 minutes at 95°C. Equal amounts of protein were loaded onto the SDS-PAGE gel. The proteins were then transferred to a PVDF membrane and stained with antibodies against pSTING (Cell Signaling, catalog 72971S), pTBK1 (Cell signaling, catalog 5483S), pIRF3 (Cell Signaling, catalog 4947S), and Actin (Cell Signaling, catalog 3700S) antibodies (1:1000 dilution) overnight at 4°C in blocking buffer, followed by staining with secondary antibodies HRP mouse anti-rabbit IgG (Santa Cruz Biotechnology, catalog sc-2357) or HRP anti-mouse IgG (Santa Cruz Biotechnology, catalog sc516102) for 1 hour at RT. Bands were visualized using a chemiluminescence-based imaging system (Bio-Rad ChemiDOC).

### Mice infection

To elucidate the role of c-di-AMP on the IFN-I response, a mice infection study was conducted following established protocols. Briefly, 4–6-week-old C3H/HeN mice of either age were organized into four groups (n=3/group). The groups were administered the following treatments: Group 1 received BSK-II medium, Group 2 received c-di-AMP (25 µg/mouse), Group 3 received *B. burgdorferi* (1×10^5 spirochetes/mouse, resuspended in BSK-II), and Group 4 received both c-di-AMP (25 µg/mouse) and *B. burgdorferi* (1×10^5 spirochetes/mouse, resuspended in BSK-II). After 1 and 7 days of infection, mice were euthanized, and the skin at the inoculation site was harvested and preserved in RNAlater within liquid nitrogen. The preserved skin tissue was subsequently used to extract RNA for qRT-PCR analysis.

### RNA Extraction and qRT-PCR

Total RNA was extracted from *B. burgdorferi* infected and mock-treated cells and tissues using TRIzol Reagent following the manufacturer’s instructions (Life Technologies, Thermo Fisher, USA). Briefly, 1 µg of total RNA was reverse transcribed using the PrimeScript 1^st^ strand cDNA Synthesis kit (Takara, catalog 6110A). The resulting cDNA was employed for qRT-PCR using PowerUp SYBR Green Master Mix (Thermo Fisher Scientific, catalog A25742) and a QuantStudio 3 Real-Time PCR thermocycler. The primers used for qRT-PCR are listed in supplemental **Table S1**. The mRNA expression levels were normalized to the housekeeping gene β-actin, and relative gene expression was determined using the ΔΔCT method (Livak & Schmittgen, 2001). Macrophages and mice tissues treated with BSK-II medium were served as the reference sample.

### Statistical Analysis

The statistical analyses were conducted using GraphPad Prism 10 (GraphPad Software, La Jolla, CA). The student’s t-test (unpaired, two-tailed) was used for all statistical comparisons between the two groups. Multiple group comparisons were performed by one-way analysis of variance (ANOVA). A value of *p*<0.05 was considered statistically significant. Error bars depict the mean ± standard deviation (S.D) from two to six biological replicates. * Indicates *p*<0.05, \*\**p*<0.01, \*\*\**p*<0.001, \*\*\*\**p*<0.0001.

## ACKNOWLEDGMENTS

Funding for this research was partly supported by Steven and Alexandra Cohen Foundation (to X. F. Yang, H. O. Sintim), NIH grants R01AI152235 (to X. F. Yang, H. O. Sintim), R01AI083640 (to X. F. Yang), and National Natural Science Foundation of China 82172319 (to M. Ye), 82072310 (to Y. Lou).

## Abbreviations

Bb: Borrelia burgdorferi
STING: Stimulator of interferon Genes
PAMP: Pathogen Associated Molecular Patterns
c-di-AMP: Cyclic di-adenosine mono-phosphate

## SUPPLEMENTAL MATERIALS

**Supplemental Figure S1.**
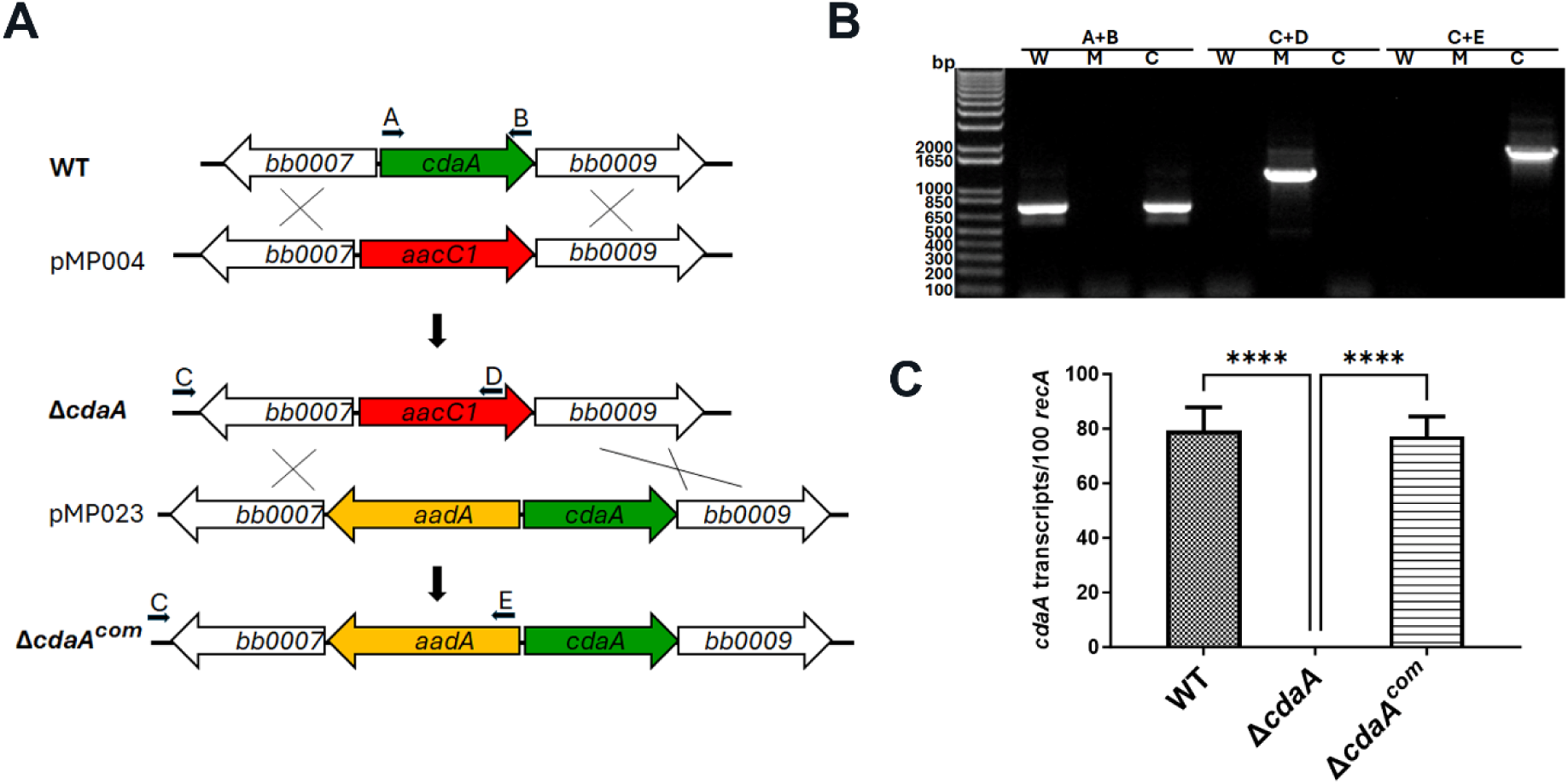
Construction of the *cdaA* mutant. (**A**) Strategy for constructing the *cdaA* mutant (Δ*cdaA*) and the complemented strain (Δ*cdaA^com^*). pMP004 and pMP023 are the suicide plasmids used for transformation and construction of the *cdaA* mutant and the complemented strain, respectively. Arrows with labels A-E indicate the positions of each primer for PCR analyses. (**B**) PCR analysis of wild-type (W), the *cdaA* mutant (M) and the Δ*cdaA* complemented strain (C) strains. The specific primer pairs used in PCR are indicated at the top. (**C**) qRT-PCR analysis of *cdaA* expression. The copy numbers of *cdaA* mRNA were normalized with 100 copies of a house-keeping gene *recA*. ***, *p* < 0.0001 using one-way ANOVA.

**Supplemental Table S1:**
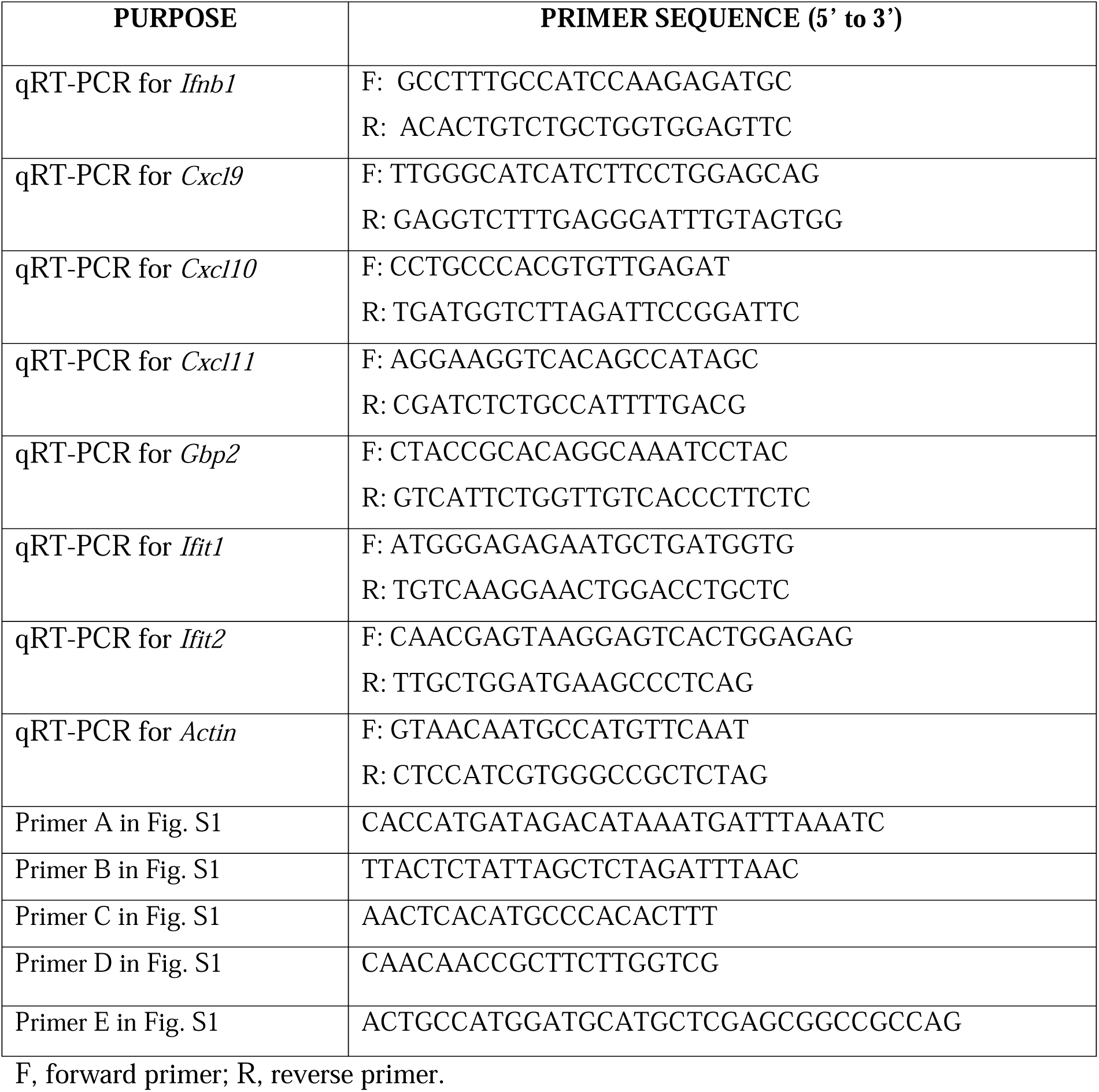
Primers used for qRT-PCR and PCR in this study.

